# A neural signature of automatic lexical access in bilinguals

**DOI:** 10.1101/2021.07.20.452909

**Authors:** Sabrina Aristei, Aliette Lochy, Bruno Rossion, Christine Schiltz

## Abstract

Bilingualism is often associated with beneficial effects on cognitive control and top-down processes. The present study aimed at bypassing these processes to assess automatic visual word recognition in bilinguals. Using fast periodic visual stimulation, we recorded frequency-tagged word-selective EEG responses in French monolinguals and late bilinguals (German native, French as second language). Words were presented centrally within rapid (10 Hz) sequences of letter strings varying in word-likeness, i.e., consonant strings, non-words, pseudo-words, while participants performed an orthogonal task. Automatic word-selective brain responses in the occipito-temporal cortex arose almost exclusively for the languages mastered by participants: two in bilinguals vs. one in monolinguals. Importantly, the amplitude of bilinguals’ responses to words within consonant strings were unaffected by the native vs. late-learnt status of the language. Furthermore, for all and only known languages, word-selective responses were reduced by embedding them in pseudo-words relative to non-words, both derived from the same language as the words. This word-likeness effect highlights the lexical nature of the recorded brain visual responses. A cross-language word-likeness effect was observed only in bilinguals and only with pseudo-words derived from the native language, indicating an experience-based tuning to language. Taken together these findings indicate that the amount of exposure to a language determines the engagement of neural resources devoted to word processing in the occipito-temporal visual cortex. We conclude that automatic lexical coding occurs at early visual processing in bilinguals and monolinguals alike, and that language exposure determines the competition strength of a language.

**Significance Statement:** Bilingualism and its possible impact on automatic processes have rarely attracted interest, contrary to bilingualism and its mutual relation with the executive functions. We assessed automatic visual word recognition in bi- and monolingual individuals while purposively bypassing executive functions. Visual brain potentials frequency tagged to words, that were flashed in rapid trains of strings with varying word-likeness degrees, exposed the automatic encoding of word-form as well as language identity at early stages of visual word processing within the occipito-temporal visual cortex. The mechanisms involved in both encoding processes reflect experience-based activity as the one characterizing tight-tuned neurons in the VWFA. Our findings provide a novel framework to understand the mechanisms behind the incredible efficiency of bilinguals in handling multiple languages.

## Introduction

Bilingualism seems to constitute a great advantage for human brains by training core functions for our everyday life such as conflict resolution (1). While benefits of bilingualism on conflict resolution abilities and the underlying neural mechanisms have been extensively investigated and sparked lively debates (1–4), the potential effects of bilingualism on automatic perceptual processes (5) remain almost unnoticed. However, bilinguals represent the natural experiment to investigate and understand how brains process conflictual information prior conflict monitoring and resolution come to aid.

In bilingual speakers, both of their known languages are jointly active and this joint activation is outside their control. Even without being physically present, known languages have the power to automatically affect verbal behavior in the currently used language and their underlying neural correlates (6–10). This uncontrollable joint language activation interferes with the verbal task at hand (1), and thus exacerbates the cognitive load linked to linguistic processing.

To assess potential effects of bilingualism on automatic perceptual processes (5), we measured brain activity during visual word recognition, a putatively automatic process in monolingual speakers. Remarkably, bilinguals show a specific pattern of results in the Stroop paradigm, known since decades to reveal automatic visual word recognition (11): compared to monolingual peers, bilinguals show smaller interference from incongruent color words, thus reduced Stroop incongruency effects (12–14). On the one hand, this lower susceptibility to conflicting input information has been taken as evidence against automatic visual word recognition in bilinguals (15), due to a slower/shallower processing of the irrelevant word. On the other hand, it has been interpreted as proof in favor of either automatic (word) processing in bilinguals, in combination with better top-down interference suppression (12–14), or alternatively, of faster disengagement of attention from distracting information (i.e., color word) and re-allocation of the newly available resources on the target input and task at hand (16).

These opposite interpretations of the very same behavioral data highlight the need to address this outstanding issue of whether and how bilingualism affects automatic visual word recognition by bypassing top-down conflict monitoring and resolution mechanisms linked to executive functions. We achieved this in the present study by measuring visual word recognition without an explicit verbal task and by varying the degree of concurrent activation within and across two languages through the word-likeness and the language membership of irrelevant items.

Specifically, we used a recently validated Fast Periodic Visual Stimulation Paradigm (FPVS) in the French language in which variable letter strings (i.e., pseudo-words, non-words or consonant strings in different conditions) appear at a rapid fixed frequency of 10 Hz, with variable words appearing periodically every five items (2 Hz) (17, 18). Thus, words appear in a rapid stream of frequent base stimuli differing in word-likeliness. Besides a strong 10 Hz (and harmonics, e.g., 20 Hz, etc.) response common to words and the respective letter strings used in a stimulation sequence (e.g., non-words), any selective (i.e., differential) reliable response to words in the sequence should elicit a 2 Hz (and harmonics, e.g., 4 Hz, 6 Hz, etc.) peak in the frequency-domain representation of the human electroencephalogram (EEG) (17). This approach has considerable advantages in terms of objectivity (i.e., neural responses are identified and quantified at an experimentally pre-defined frequency) and sensitivity (i.e., high signal-to-noise ratio), making it ideal to probe automatic visual word recognition processes (19, 20). In the current paradigm, a fixation cross is presented centrally and concurrently with the letter strings, requiring observers to detect its (non-periodic) random color changes. This task is thus fully orthogonal to word recognition, with verbal stimuli and language processing being totally irrelevant.

Here, critically, to investigate automatic visual word recognition in bilinguals for the first time, we measured brain responses to French and German words in German-French bilinguals and compared them to French native speakers, under three conditions with varying degrees of word-likeness. Base stimuli consisted of the same consonant strings (e.g., XWZRT) used for both groups of speakers, non-sense letter strings including consonants and vowels referred to as non-words (e.g., LMUBE), and letter strings orthographically plausible and thus similar to real words, referred to as pseudo-words (e.g., MUBEL similar to the German word MÖBEL, furniture pieces). Non-words and pseudo-words shared the same letters with the deviant words they were derived from and were therefore language-specific (from German words: BLUME, flower in German, non-word – LMUBE, and pseudo-word – MUBEL). Therefore, we had a set of German-derived and French-derived non-words and pseudo-words, that we refer to as German/French non-words and pseudo-words per convention, even though only pseudo-words have the merit to be categorized according to a language.

Assuming automatic visual word recognition, such that abstract word representation is automatically accessed in monolinguals and bilinguals alike, the amplitude of a word-selective neural response should be unaffected by base stimuli that do not compete for lexical processing resources. Therefore, word- selective brain responses should be of maximal amplitude in the consonant string condition, irrespective of the word language. Yet, for French monolinguals, no word-selective activity is expected in response to German words as these do not have a word status and hence, no abstract representation. Furthermore, in both groups, word- selective brain responses should display a left-lateralized posterior distribution, as in previous FPVS-studies in monolinguals (17, 18), and in line with the role of the left ventral occipito-temporal cortex (VOTC) in visual word recognition (21, 22).

Likewise, when using non-words as base stimuli, automaticity of visual word recognition in both of bilingual’s languages should induce responses of similar amplitude to German and French words, independently of the language of derivation of the non-words. Any difference related to the language of the non-words would reflect the involvement of sources that contribute to sub-lexical processes, for instance, because words and non-words shared all letters in the within- and not in the cross-language condition. Again, for French monolinguals, only French words should produce word-selective neural responses.

Since pseudo-words activate partially matching words stored in the long-term memory, they strongly compete with resources deployed for the recognition of the deviant word stimuli (23–25) and we should consequently observe reduced neural responses to words within pseudo-words, compared to the other conditions (17). Hereafter we will refer to this effect as the word-likeness effect, using it as an index of lexical competition.

Furthermore, as form similarity is an organization principle of the lexicon (26), higher competition should occur between pseudo-words and words from within the same language. Hence, in bilinguals within-language pseudo-words should reduce the word-selective brain response for both of their languages even further than cross-language pseudo-words. As French monolinguals do not have a lexicon for German words, we again expected only a reduction to French (but not to German) words embedded in pseudo-words. Furthermore, German pseudo-words should act similarly to non-words for the French monolinguals.

Our findings will inform on automatic word recognition in bilinguals compared to monolinguals, revealing how visually presented words in both of a bilingual’s languages are being processed independently of executive control.

## Results

### Automatic word-selective neural responses only for known languages

EEG responses to words within consonant strings provide an index of sensitivity to lexical inputs when competition is minimal and equal for both sets. For words within consonant strings, z-score topographies of the grand means revealed significant responses at the first four harmonics (2 Hz, 4 Hz, 6 Hz, 8 Hz; z-scores > 1.64, p <.05) with an occipito-temporal left lateralized distribution (Figure 2A and B). This left occipito-temporal topography accords with the typical language lateralization in the literature (27, 28), and with the topography of FPVS-induced brain responses to visual words in monolingual studies (17, 29).

**Figure 1.**
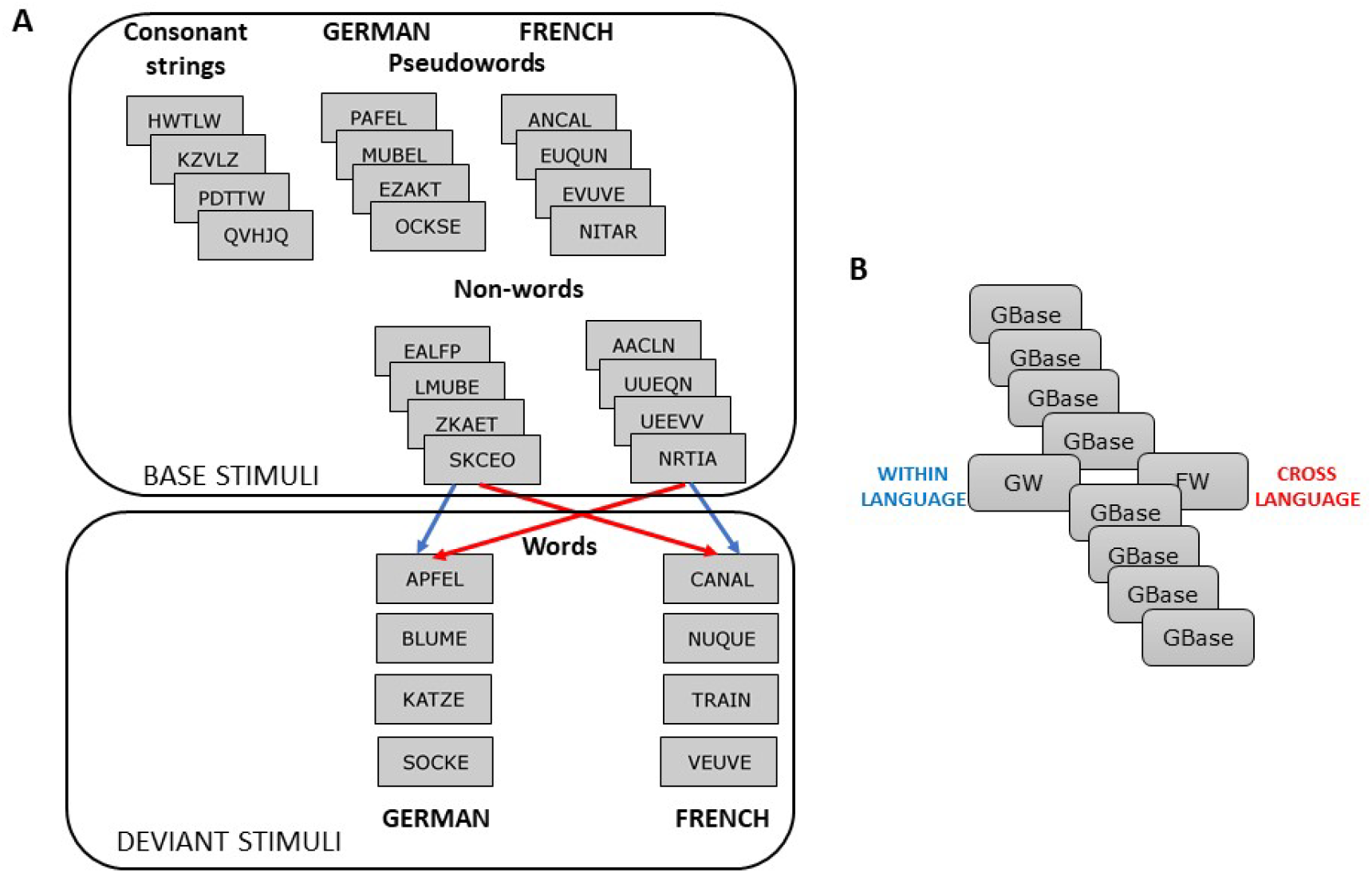
Material and trial design. (A) Examples of the stimulus set for each base and deviant condition and (B) trial scheme.

**Figure 2.**
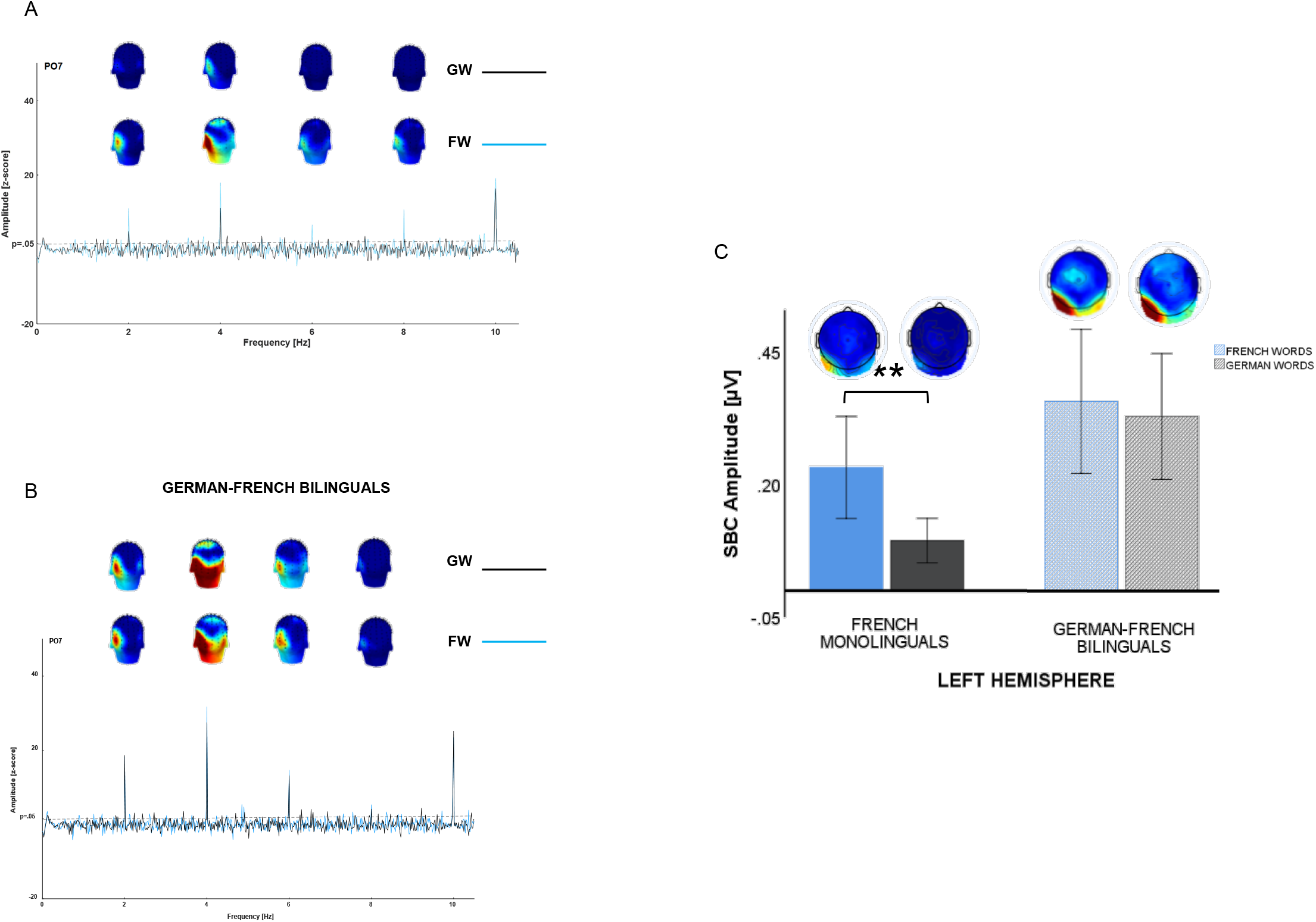
Grand average of the brain responses to French (blue line) and German (black line) words (FW, GW) within Consonant Strings. (A) Plots of z-scores with the respective significance maps for the first four harmonics (2-8 Hz) for French speakers and (B) German-French bilinguals. (C) Plots of the baseline corrected amplitude summed across the first four harmonics (SBC) for the left ROI (signal averaged across electrodes within the left cluster).

The amplitude of the neural response to French and German words depended on the participants’ profile and language knowledge. In French monolinguals, robust word- selective responses arose in response to French words, and exclusively over the left occipitotemporal region (left lateralized response, French words: 0.23 μV; German words: 0.09 μV; see figure 2C). In German-French bilinguals to the contrary, a brain response of roughly equal amplitude emerged for both French and German words (left hemisphere: 0.36 μV and 0.33 μV, respectively; see figure 2C).

The difference in word-selective neural activity between mono- and bilinguals was confirmed with a repeated measures ANOVA (Electrodes × Hemisphere × Word Language as within-participants factor, and Language Group as between-participants factor), which yielded a highly significant interaction between language of word and language group, F(1,41) = 10.80, p = .002, ηp2 = .209. Overall, word-selective brain responses in bilinguals were of larger amplitude (group main effect: F(1,41) = 8.49, p = .006, ηp2 = .172) and had a more bilateral scalp distribution than in French monolinguals (hemisphere × language group interaction, F(1,41) = 8.82, p = .005, ηp2= .177; see scalp maps in Fig. 2C).

Overall, the pattern of results show that robust word-selective responses (as compared to consonants strings) emerge specifically for known languages, and converge with the hypothesis of automatic word-selective neural responses in both mono- and bilingual participants

### Automatic within-language competition for visual neural resources

In contrast to consonant strings, responses to words within non-words and pseudo-words from the same language provide a direct test of how sensitive word-responses are to lexical competition. Since the latter within-language base stimuli do not differ from words in terms of shared low-level visual features or letters, only competition for lexical resources can determine different word-selective responses in the two conditions. The word-likeness effect is computed by subtracting the amplitude of word-selective responses in pseudo-words from those to words embedded in non-words.

The effects of competition on the neural responses to words over bilateral occipito-temporal regions was confirmed by a word-likeness main effect, F(1,41) = 11.78, p = .001, ηp2= .223, reflecting a strongly reduced response to words within pseudo-words compared to words presented in the non-word context (see Figure 3B and 3C) alike previous studies in monolinguals (Lochy et al., 2015). Most importantly, participants’ language profile had an impact on word-selective responses (word-likeness × language group: F(1,41) = 6.49, p = .015, ηp2= .137). Subsequent separate ANOVAs for mono- and bilinguals clarify the differences between the two groups.

**Figure 3.**
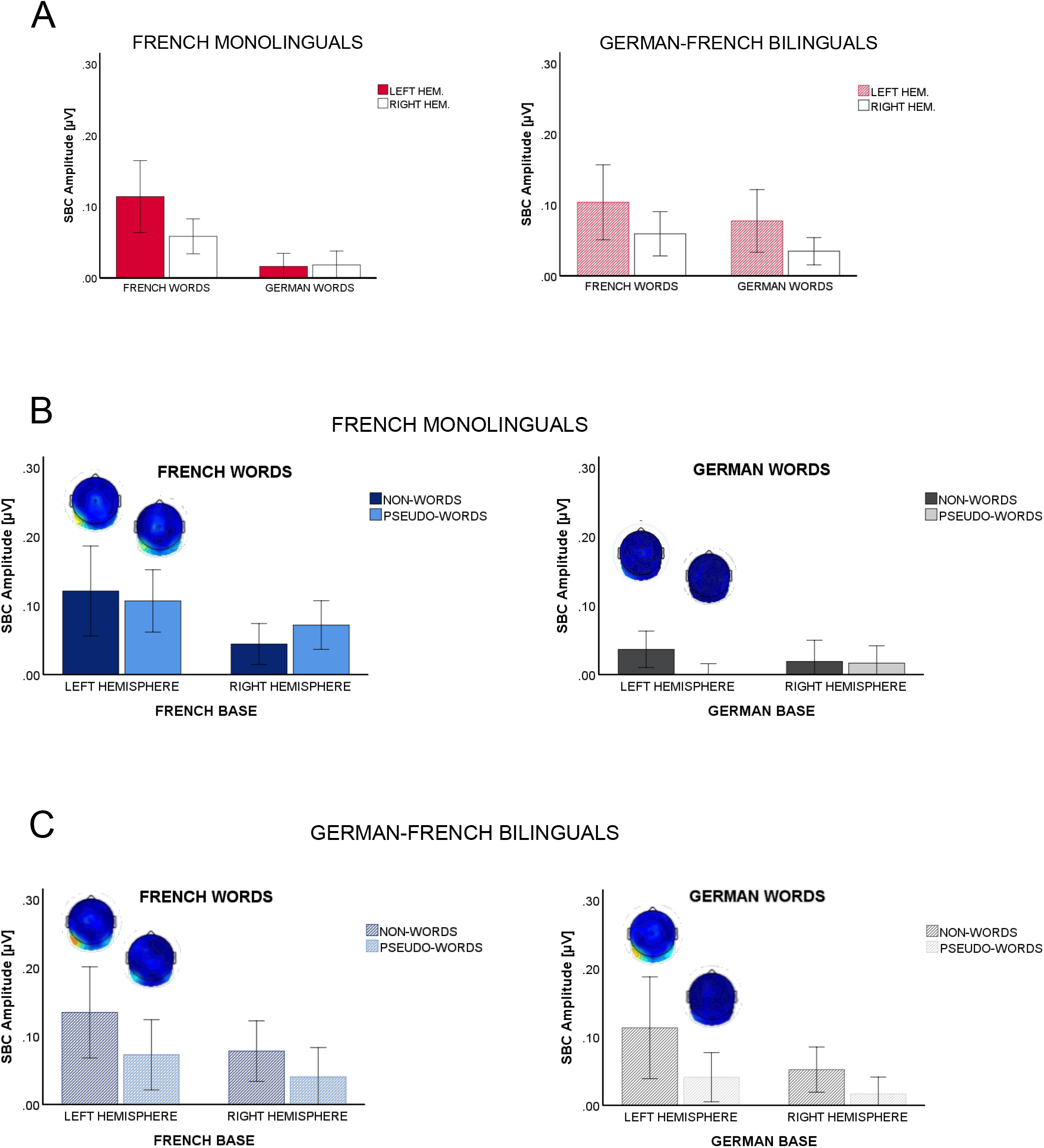
Within-language effects. The baseline-corrected amplitudes summed across the first four harmonics (SBC) are plotted for the word-sensitive response to French and German words in the within-language conditions. Plots display the difference in the amplitude of the brain response to words (within a sequence of non-words or pseudo-words) as a function of the participants language profile (A), and as a function of word-likeness of the base stimuli for French monolinguals (B) and German-French bilinguals (C) with the respective scalp topographies.

In French monolinguals, robust word-selective responses were again elicited in response to French words, while responses to German words were almost non-existent (language: F(1,19) = 33.03, p < .001, ηp2= .635; across conditions, LH - mean amplitude for French words: 0.114 μV; for German words: 0.016 μV; see Figure 3A, left plot). Word-likeness affected left lateralized word-selective responses (word-likeness × hemisphere: F(1,19) = 5.82, p = .026, ηp2= .234; LH – word-likeness: F(1,19) = 4.5, p = .047, ηp2= .192; RH: F(1,19) = 1.49, p = .24, ηp2= .073; see Figure 2B).

In German-French bilinguals, word-selective responses were also significantly reduced within the pseudo-word context by comparison to the non-word context, this time in both languages (word-likeness effect: F(1,22) = 12.87, p = .002, ηp2= .369; see Figure 2C). Interestingly, the word-likeness effect was stronger than in monolinguals irrespective of language (response amplitude for French words in pseudowords: monolinguals, 0.114 μV; bilinguals, 0.062 μV).

Language main effects, F(1,22) = 5.67, p = .026, ηp2= .205, revealed a larger amplitude for French than German words across conditions and hemispheres. They were driven by the almost doubled amplitude reduction on word responses produced by German than French pseudo-words (see Figure 3C). Additionally, while being left lateralized for monolinguals, the word-likeness effects presented no lateralization in bilinguals (word-likeness × hemisphere: F(1,22) = 1.97, p = .17, ηp2= .082; see Figure 3C).

In sum, our measure of automatic selective responses to printed words consistently reflected the participants’ language profile, with word-selective responses and word-likeness effects reflecting language knowledge. The drop in the response amplitude confirmed automatic competition for lexical resources, which had an impact on automatic word-selective responses to known language(s), similarly in bilinguals and monolinguals.

### Differential cross-language competition for native and late learnt language

Results in the cross-language conditions, using base stimuli derived from words of the other language, allowed to assess how sensitive word-responses are to competition across languages. As for the within-language conditions, non-words and pseudo-words contained the same letters. Even if these letters differed from the words’ letters, both base conditions visually differed to the same degree from the word deviants, which excluded the contribution of mere visual and pre-lexical factors to differences between conditions. This assured that word-likeness effects reflect concurrent cognitive, high-level processes.

As for the within-language comparisons, we analyzed the neural activity over occipito-temporal regions in response to words embedded in non-words and pseudo-words, this time of the other language (i.e. German words within French base stimuli and vice versa for French words). Bilingualism and input language affected word-selective responses and modulated the strength of competition exerted by pseudo-words (language × language group, F(1,41) = 23.87, p < .001, ηp2= .368; language × word-likeness, F(1,41) = 4.81, p = .034, ηp2= .105; language × word-likeness × language group: F(1,41) = 3.34, p = .07, ηp2= .075).

In French monolinguals strong word-selective responses were elicited again for the known language, French. Once more word-selective response to German words was very weak, irrespective of the base condition (see Figure 4A and B), corroborating the idea that word-selective responses reflected the activity of lexical populations of neurons. Maximal amplitudes were found over the left hemisphere (language: F(1,19) = 15.95, p = .001, ηp2= .456; hemisphere × language: F(1,19) = 6.65, p = .018, ηp2= .259 mean amplitudes LH – French words: .134 μV; German words: .057 μV).

**Figure 4.**
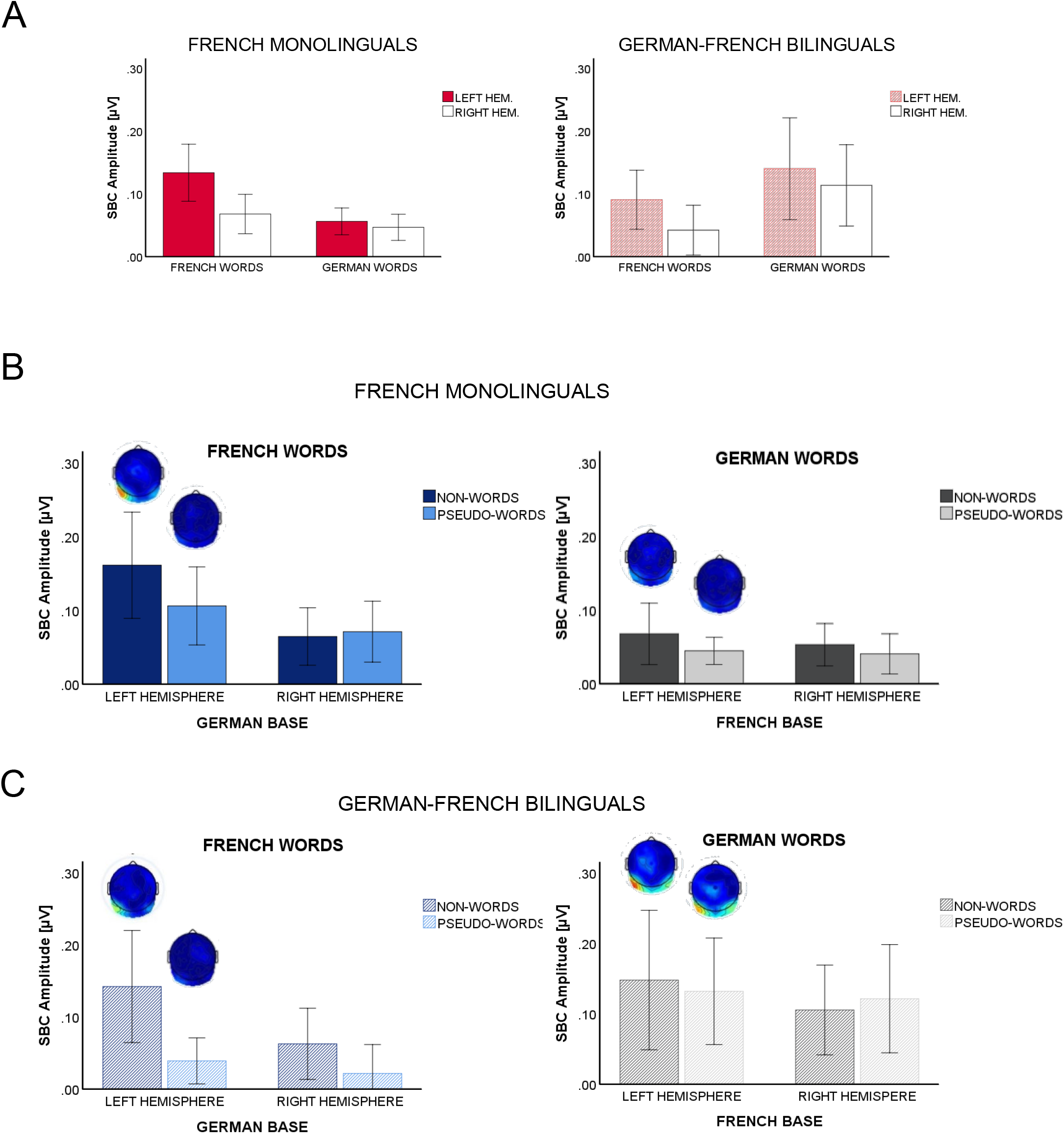
Cross-language effects. The baseline-corrected amplitudes summed across the first four harmonics (SBC) are plotted for the word-sensitive response to French and German words in the cross-language conditions. Plots display the difference in the amplitude of the brain response as a function of (A) the participants language profile, and as a function of word-likeness of the base stimuli for (B) French monolinguals and (C) German-French bilinguals with the respective scalp topographies.

As German words are treated as non-words, that is, unknown letter strings, the word-likeness of German base stimuli did not significantly affect word-selective responses to French words (word-likeness effects: F(1,19) = 3.09, p = .09, ηp2= .140; language × word-likeness: F < 1; hemisphere × word-likeness: F(1,19) = 2.83, p = .11, ηp2= .130; hemisphere × language × word-likeness: F(1,19) = 1.77, p = .2, ηp2= .085). Thus, in monolinguals letter strings mimicking the orthography of an unknown language did not affect the word-responses to the known native language.

In German-French bilinguals (see Figure 4C), the amount of competition induced by pseudo-words relative to non-words depended on the input language, i.e. native vs. late learnt language (language × word-likeness: F(1,22) = 10.56, p = .004, ηp2= .324; word-likeness effects: F(1,22) = 5.09, p = .034, ηp2= .188). Separate ANOVAs for the two languages (electrodes × hemisphere × word-likeness) revealed that word-selective responses were reduced in pseudo-words by comparison to non-words only in the presence of a cross-language competition from the native language, that is, when French words appeared within German pseudo-words (word-likeness effects – French words/German base: F(1,22) = 12.24, p = .002, ηp2= .357) and not vice versa (German words/French base: F(1,22) < 1).

Furthermore, the word-selective responses to French words were left lateralized (hemisphere × word-likeness: F(1,22) = 6.87, p = .016, ηp2=.238), contrary to the within-language conditions. Similarly, the topography of the word-selective responses to German words showed bilateral responses (hemisphere × word-likeness: F(1,22) < 1), even more strongly than in the within-language conditions.

In sum, in bilinguals, pseudo-words had a cross-language impact only from the native onto the late-learnt language, and not vice versa. That is, while German pseudo-words reduced the brain response to French words, French-derived pseudo-words did not reduce word-selective responses to German words. Moreover, in French monolinguals, pre-lexical characteristics of the base stimuli (i.e., pseudo-words vs. non-words) from the unknown language German had no significant effect on the brain responses to French words. Likewise, word-selective responses to German words were very weak compared to French words, indicating that brain responses to words were driven by the lexical status of the input stimuli, rather than low-level visual or pre-lexical stimulus characteristics. Thus, input lexicality and input language together determined the deployment of resources for word recognition.

## Discussion

By capitalizing on the sensitivity of FPVS-induced brain responses to the contrast between the deviant and base stimuli, we found evidence for the automatic activation of bilinguals’ two known languages when none of them was in use. Different levels of competition contrast were tested by presenting words in rapid succession intermingled with either consonant strings, orthographically illegal letter strings (non-words), or orthographically legal letter strings (pseudo-words). Evidence in favor of automatic visual word recognition was provided by three findings. First, robust word-selective neural responses arose specifically for the known language or languages (in bilinguals). Second, word-selective responses were unaffected by competition for resources engaged in pre-lexical processes (i.e., words vs. implausible letter strings) but were modulated by the level of lexicality of the base stimuli (i.e., nonwords or pseudowords). Third, cross-language competition effects occurred only for bilinguals and was only induced by the most trained language (i.e., the native language).

An EEG signature of automatic bilingual word recognition. The first level of assessment aimed at setting a baseline of the bilinguals’ ability to automatically discriminate words in the two known languages with the broadest contrast, i.e., consonant strings. It conveyed a clear electrophysiological signature of automatic bilingual word recognition. Indeed, words from both native and late learnt languages (German and French, respectively) elicited word-selective responses of equivalent amplitude. In contrast and as predicted, for French monolinguals a robust response to words arose in the one known language (French).

At this discrimination level (words vs. consonant strings), the pattern we found in bilinguals suggests automatic access to a high-level word representation for all known languages, irrespective of properties often found to affect language processing such as language proficiency, age of acquisition and frequency of use (30). Apparently, the determinant factor driving the strength of word-selective brain responses was language knowledge (known/unknown), suggesting that they likely reflect the activity of category-selective units (31). Against consonant strings, word discrimination could alternatively be driven by low-level visual differences because the base and word stimuli contain different letters, or by differences pertaining to single letter or letter clusters as only word stimuli contained consonant-vowel structures (32, 33). However, if implicit word discrimination were based only on visual dissimilarity or letter discrimination, then responses to words should have been insensitive to language membership and to the language profile of the speakers. To the contrary, our results showed a difference in the response pattern between mono- and bilingual speakers, which excludes that responses to words in this condition reflect merely a visual dissimilarity or a letter-detection mechanism.

The deterministic nature of the response to words driven by the mastery of the presented language allows for a reliable distinction between monolingual and bilingual lexical coding. Therefore, this response may constitute a reliable brain marker of multilingualism.

Lexical modulation of word-selective responses in within-language conditions. The automaticity of visual word recognition was further evaluated by using word-like letter strings, i.e., pseudo-words as base stimuli, in which words were periodically embedded. Because of their high similarity, the discrimination between pseudo-words and real words is thought to require the engagement of partially overlapping neuron assemblies (31), and as a consequence, to result in competition for neuronal resources (34, 35).

As expected, in the within-language conditions, when both non-words and pseudo-words shared the same letters with words, word-selective brain responses were lower in the pseudo-words relative to the non-words condition, i.e. the word-likeness effect (17). This effect is interpreted as an increase of competition for neural resources. In bilinguals, the drop in the neural response occurred for both languages, with word-selective responses in native and late-learnt language similarly affected by pseudo-words derived from the same language. In monolinguals, the word-likeness effect occurred only for the known language, French. As no word-selective response arose for German words, no amplitude reduction could be measured. Thus, competition between words and pseudo-words for neural representation occurred for stimuli from all languages mastered by the participants. The observed difference between monolingual and bilingual participants in the non-word and pseudo-word conditions converges with the pattern of results in the consonant string condition. The language-specific sensitivity of the brain responses to words supports a category-selectivity of their neuronal source, which depends on having or not extensively experienced a given language.

Additionally, word similarity of the base stimuli drives the competition for word-processing representations, diming the detectability of the word-selective brain response when words are embedded in more (i.e. pseudo-words) compared to less similar (i.e. non-words) stimuli. At a first glance, the discreteness of the word-selective mechanism observed in the consonant-string condition might seem incompatible with the sensitivity to the word-likeness of base stimuli. However, intracerebral FPVS recordings of monolingual word processing reported word-responsive contacts that were and were not affected by the word-likeness of the base stimuli within the same region in the left fusiform gyrus (18). Thus, category-selectivity and word-likeness modulations of the brain responses to words likely reflect the activity of these two partially overlapping neural populations. The intracerebral findings of monolingual word processing and our findings in bilinguals point to two distinct neural mechanisms reported within the specific structure of the left fusiform gyrus, the Visual Word Form Area (VWFA) (36): a tight tuning to real words and a broad tuning to pseudo-words (31, 37) (for a similar mechanism in face recognition see (38)).

The response selectivity for known/unknown languages we observed across conditions resembles the behavior of word-selective tightly-tuned neurons within the VWFA, which are invariant to different degrees of orthographic similarity of adjacent items. By contrast, the sensitivity to the word-likeness of base stimuli resembles the behavior of broadly-tuned neurons, which respond also to untrained letter strings such as pseudo-words, and in contrast to the selectivity of the tight tuned cells, their activation is proportional to the input word similarity (31). These two mechanisms may take place in partly overlapping neural populations, and when they are activated simultaneously and to a similar degree, brain responses to the word-like letter strings (broad tuning) compete with and consequently dim the relative amplitude of word-selective responses (tight tuning) as recorded on the scalp. Therefore, scalp recorded word-selective visual evoked potentials likely reflect the combined activation of these two populations of tight and broad tuned neurons. The occipito-temporal topography of the effects observed in the present study further support the view that visual brain responses specific to words of known languages and the word-likeness effect reflect processing of these different neural populations within the VWFA. Language dominance as determinant of automatic cross-language competition. The cross-language conditions revealed an even more refined scenario, by providing compelling evidence that the recorded brain responses reflected not only the overall learnt/unlearnt status of a language, but also the differential mastery of languages. Again, in bilinguals, words in both known languages evoked a word-selective brain response, whereas in French monolinguals a robust response was observed to the known language, i.e., French words. Likewise, a cross-language word-likeness effect arose exclusively in bilinguals. The effect consisted in a drop of the word-selective response to French words within German pseudo-words relative to German non-words. In contrast in monolinguals, responses to French words were largely unaffected by the word-likeness of base stimuli in the unknown language (i.e. German pseudo-words vs. German non-words).

Furthermore, in bilinguals the German base stimuli induced word-likeness effects both in the within and the cross-language conditions, whereas French base stimuli entailed word-likeness effects exclusively in the within-language condition. That is, in the cross-language condition word-selective visual brain responses to German words were equally strong irrespective of the word-likeness of the French base stimuli in which they were embedded. The brain responses evoked by French pseudo-words thus seemed too weak to compete with and to dim the tight-tuned neural response to German words. Thus, in the suggested scenario where the response to words recorded on the skull reflects the combined visual resources engaged for word (tight tuned neurons) and pseudo-word (broad tuned neurons) processing, the efficacy of broad tuned neurons seems to depend as well on the experience with a language and language-specific properties. In this case, more experience with a language would entail more efficient – stronger or faster –activation of broad tuned neurons in response to pseudo-words, i.e., letter strings that closely follow language-specific orthographic rules.

Hence, besides evidence for automatic brain responses to words of known languages, bilinguals’ brains revealed differential experience-based language activation encoded within the occipito-temporal visual cortex. This is evident in the distinct effects of base stimuli, as native German pseudo-words were powerful competitors irrespective of the language words belonged to, while later learned French pseudo-words were too weak competitors to produce a response drop in German word-selective responses.

Alternatively, rather than the amount of exposure to a language, distinct degrees of language transparency may determine the difference in competition between languages, with more efficient activation of broadly tuned neurons for the most transparent language (i.e. German). Both accounts are compatible with the co-existence of tight-tuned word selective neurons and broader tuning to pseudo-words within the VWFA (31). However, the whole pattern of results and the occipito-temporal topography of word-responses, mostly left lateralized, strongly accords with the neuro-functional localization of the two anatomically overlapping mechanisms in the VWFA. Conclusion. In sum, FPVS and neural visual responses revealed automatic access to visual word forms of all known languages. Despite the fact that input letter strings were totally irrelevant for the task, neuronal populations automatically responded selectively to their preferred stimulus category – words, in any of the known languages. Similarly, the amount of experience with a given language determined how strongly word-like stimuli (within and cross-language) compete with words for visual representations, as reflected in the drop of the brain responses to words (i.e., word-likeness effect). We advocate that both the automatic activation of all known languages, and the language-dependent competition reflect distinct experience-based mechanisms, in line with two neural mechanism within the VWFA, a tight-tuning to words and a broad-tuning to word-like letter strings (31, 37). Therefore, our findings indicate that to some extent language is, alike word-form, automatically encoded at early stages of visual word processing within the occipito-temporal visual cortex. This contrasts with previous attempts to locate assumed language identity processing in high-level frontal areas (39), which are typically associated with executive functions, those we purposively bypassed in our study. Beyond these new insights on automatic visual word recognition in bilinguals, frequency-tagged word-selective brain responses can serve as a signature of how many and which languages are in a person’s brain, thus providing a brain marker of multilingualism.

## Materials and Methods

### Participants

We tested forty-four participants, 24 German-French late bilinguals and 20 French native speakers (age range: 19-29 years old) with normal or corrected-to-normal vision. One German participant was excluded from the analysis because of a failure in the stimulation system. All participants had a monolingual family background and schooling (only German, only French respectively). German-French bilinguals were late learners having acquired French between 10 and 13 years old. No participants had a reported history of neurological or psychiatric disorder. All participants were studying or working in Luxembourg, that is, they were exposed to the same multilingual environment characterized by the three official languages in the country which are Luxembourgish, German and French. Experimental procedures were conformed to the World Medical Association Declaration of Helsinki and the local institutional ethic committee approved the study.

Language Assessments. Before the experimental session, participants carried out an online test, which assessed their proficiency in both German and French by standardized tests of grammatical knowledge, vocabulary and reading comprehension (www.transparent.com). German-French bilinguals scored between 97% and 100% for German, and between 60% and 88% for French. This corresponds to a medium-high proficiency, except for one participant who scored 52%. French native speakers scored between 87% and 97% for their native language, French, and between 0% and 31% for German, confirming no formal knowledge of this language (two participants had higher scores 49% and 69%).

### Stimuli

We selected 30 high-frequent German nouns (e.g., “KATZE”), and thirty high-frequent French nouns (e.g., “ARBRE”). All words were five letters long and did not include foreign words, accented words, or translation equivalents. Descriptives of linguistic variables were determined with Word Gen (CELEX database) and are reported in Table 1.

**Table 1.** Description of lexical and sub-lexical indicators for the German and French words. Summed neighbors (N)-density is equal to the number of neighbors (i.e., words sharing the same letters but one) across languages.

|  | Lemma Frequency | N-density | summed N-density | Bigram Frequency |
| --- | --- | --- | --- | --- |
| GERMAN WORDs | 1.37 | 2.93 | 5.63 | 13767.00 |
| FRENCH WORDs | 1.78 | 1.60 | 2.13 | 12049.00 |
|  | -0.42* | 1.33* | 3.5* | 1718 |

\* p-value < .05

To build non-lexical stimuli, we re-arranged each German and French word into corresponding illegal non-words (e.g., German: ZKAET; French: RRBAE), and pseudo-words (e.g., German: EZAKT; French: ERBRA), for a total of 30 non-words and 30 pseudo-words. In addition, 30 five-letter long consonant strings (e.g., SVZQC) were generated with Word Gen software (40).

Words and non-lexical stimuli were presented in sequences. Within each sequence, words appeared at regular intervals, that is periodically, every fifth stimulus (ratio 1/5). Likewise, we refer to non-lexical strings (consonant strings, non-words and pseudo-words) as base stimuli. Within each sequence word deviants were repeated three times and base stimuli 16 times, for a total of 90 word deviants and 480 base stimuli, amounting to 600 items per sequence.

Sequences were blocked for the different base stimuli and word language, that is, German- and French- derived base stimuli were always presented in separate sequences, and so were German and French words. This led to the following 10 conditions: The first two conditions presented consonant strings as base stimuli and words of each language as word deviants (i.e., CSN-FW and CSN-GW). The 4 within-language conditions consisted of FW or GW presented among same-language NW and PW (FNW-FW; FPW-FW; GNW-GW and GPW-GW). The 4 across language conditions consisted of FW and GW presented among the base stimuli derived from the other language (Figure 1).

This entailed that German word deviants (e.g., GW: KATZE), were presented in two within-language sequences, which consisted of their corresponding non-words (GNW-sequence: e.g., ZKAET) and their corresponding pseudo-words (GPW-sequence: e.g., EZAKT). They were also presented in two cross-language sequences, which consisted of French non-words and French pseudo-words (FNW-sequence: e.g., RRBAE; FPW-sequence: e.g., ERBRA). For French word deviants (e.g., ARBRE), within-language sequences consisted respectively of French non-words (FNW-sequence: e.g., RRBAE) and French pseudo-words (FPW-sequence: e.g., ERBRA). Cross-language sequences consisted of German non-words (GNW-sequence: e.g., ZKAET) and German pseudo-words (GPW-sequence: e.g., EZAKT).

Each specific deviant-base sequence was repeated three times. The order of the first two blocks was fixed. German-French bilinguals saw German words within French pseudo-words (all three repetitions) first, followed by French words within German pseudo-words. For French speakers the order was inverted. This was to avoid possible pseudo-word effects from the weakest language (non-native) which might be due to previous exposure to words in that same language. For all other conditions, block and trial order was randomized on an individual basis for each participant and each repetition.

### Procedure

Upon arrival in the laboratory, participants were engaged in conversations exclusively in English throughout the whole experiment, in order to avoid priming any of the tested languages. Presentation of 10 items per second on a 100 Hz resolution screen (actual resolution was 99.992 Hz) produced a 10 Hz stimulation frequency (i.e., base rate) and a stimulus duration of 100 ms (10 screen cycles). The word deviants were thus presented at a 2 Hz rate (i.e., 10/5; deviant frequency). Visual stimulation followed a periodic sine wave of the stimulus physical contrast, each string reaching full contrast halfway through a stimulus cycle (approximately 50 ms after onset) and going back to full transparency at the end of the cycle, i.e., 100 ms after onset (17, 19, 41). Stimuli were thus visible for approximately 60 ms (minimum contrast 35%). All strings were presented centrally against a light-grey background. Stimulation sequences started with a central fixation cross in blue color. After a time interval between 2-5 s elapsed (randomly determined for each trial), a 2 s fade-in period followed and thereafter the 60 s stimulation (600 stimuli – including 120 deviants – × 100 ms duration) unfolded. At the end of the 60 s, stimulation gradually faded-out within 2 s. During the stimulation, the fixation cross was always on the screen. Participants were instructed to keep their gaze on it and to respond to color changes in the fixation cross by pressing the space bar with their right hand as fast as possible. Color randomly changed very rapidly (from blue to red (colors were isoluminant) 15 times per sequence. In total the experiment lasted 1 hour, breaks excluded. Participants could rest during self-paced breaks at the end of each 1-minute stimulation period.

### EEG acquisition

The EEG data were recorded from 128 encephalic active electrodes arranged according to the 10-20 coordinate system (https://www.biosemi.com/headcap.html), and using a BioSemi ActiveTwo equipment (BioSemi B.V., Amsterdam, Netherlands). The signal was digitized at a sampling rate of 2.048 KHz and referenced to the common mode sense (CMS). Offsets of the electrodes did not exceed 30 mV. Prior to pre-processing the signal was downsampled to 512 Hz. EEG preprocessing and analysis were performed using Letswave 6, an open source toolbox running over MATLAB (MathWorks, USA; www.nocions.org/letswave/).

### EEG Preprocessing and analyses

EEG frequencies below 0.1 Hz and above 70Hz were reduced by means of a 4th order bandpass Butterworth filter. The filtered signal was re-referenced to a common average of all channels and segmented into 60-s epochs (from the end of the fade-in to the start of the fade-out phase) for all conditions. We used ICA-decomposition of the segmented signal to identify components that, based on their topography, were unambiguously related to eye movements and other extra-cerebral activity (e.g., muscular tension), and subtracted them from the data (42). After artifact corrections, for each condition the individually recorded sequences were averaged across the corresponding three repetitions (3 × 60s epochs), in order to increase SNR. For analysis in the frequency domain, the averaged data were Fourier-transformed (FT) to their frequency amplitude spectrum with a 0.0167 Hz (1/60 s) resolution.

We then calculated z-scores on the grand mean of the FT-data for each condition separately and relative to the twenty surrounding frequency bins (+/− 0.167 Hz; (43)). We selected posterior electrodes that exceeded a threshold of 1.685 (alpha level = 0.05) at four consecutive harmonics. In line with previous FPVS studies of word recognition in monolinguals (17, 29) word-selective responses were significant at the first four harmonics (2 Hz, 4 Hz, 6 Hz, 8 Hz) over occipito-temporal electrodes (from the most anterior: P5, P7, P9, PPO5, PO7, PO9, POI1, O1, POO5 and right homologues).

Statistical analyses were conducted on baseline-corrected amplitudes, which we calculated by subtracting the average amplitude of the twenty bins adjacent to the harmonic of interest (19, 44) and, subsequently summed over the four selected harmonics (43). For the analyses using consonant strings as base stimuli, ANOVAs with repeated measures included electrodes (9 levels), hemisphere (2 levels), word language (2 level). In the analyses using non-words and pseudo-words as base stimuli we added word-likeness of the base stimuli as factor (2 levels). In all analyses we included group (monolingual, bilingual) as a between participant factor. Where the effects of interest interacted with language group, we conducted separate ANOVAs for the two groups, keeping all other factors as in the overall ANOVA (electrodes × hemisphere × word language and word-likeness in analysis using non-words and pseudo-words as base stimuli). Similarly, we conducted separate ANOVAs for the two hemispheres in order to decompose interactions arisen between the effects of interest and the hemisphere factor.

## Acknowledgments

We thank Julie Lang and Tonie Schweich for assisting during data collection.

## Notes

### Competing Interest Statement

The authors have declared no competing interest.

